# Origin, evolution, and distribution of the molecular machinery for biosynthesis of sialylated lipooligosaccharide structures in *Campylobacter coli*

**DOI:** 10.1101/225771

**Authors:** Alejandra Culebro, Miguel P. Machado, João André Carriço, Mirko Rossi

## Abstract

*Campylobacter jejuni* and *Campylobacter coli* are the most common cause of bacterial gastroenteritis worldwide. Additionally, *C. jejuni* is the most common bacterial etiological agent in the autoimmune Guillain-Barré syndrome (GBS). Ganglioside mimicry by *C. jejuni* lipooligosaccharide (LOS) is the triggering factor of the disease. LOS-associated genes involved in the synthesis *(neuABC)* and transfer of sialic acid (sialyltranferases) are essential in *C. jejuni* to synthesize ganglioside-like LOS. Therefore these genes have been identified as GBS markers. So far, scarce genetic evidence supports *C. coli* as a GBS causative agent despite being isolated from GBS patients. Here we show that genes putatively involved in sialic acid transfer are widely distributed in the *C. coli* population. Evidence found herein suggests that a small group of *C. coli* strains are very likely to express ganglioside mimics, implying that *C. coli* can potentially trigger GBS. *C. coli* also presents a larger repertoire of sialyltransferases than *C. jejuni* and loss of functions of some those LOS-associated genes has happened during adaptation to agriculture niche. Nevertheless, the activity of these sialyltransferases and their role in shaping *C. coli* population is yet to be explored.

## INTRODUCTION

Glycan mimicry is a strategy utilized by pathogens to evade detection by the host innate immune system^1,2^. *Campylobacter jejuni*, the most commonly reported cause of gastroenteritis in the world, boasts a large repertoire of human glycans^3^. Molecular mimicry between sialylated *C. jejuni* lipooligosaccharides (LOS) and gangliosides may result in the onset of Guillain-Barré syndrome (GBS)^4,5^; an autoimmune acute progressive polyradiculoneuropathy with approximately 5% mortality rate^6^. To express ganglioside-like LOS^7–9^, *C. jejuni* synthesizes cytidine-5′-monophospho-*N*-acetylneuraminic acid (CMP-Neu5Ac) from uridine-5′-diphosphate-*N*-acetylglucosamine (UDP-GlcNAc) by the consecutive actions of an *N*-acetylglucosamine-6-phosphate 2-epimerase (NeuC), a sialic acid synthase (NeuB), and a CMP-Neu5Ac synthase (NeuA)^10^. Then, CMP-Neu5Ac is transferred by either of the LOS associated sialyltransferases; CstII (α2,3/8-sialyltransferase) or CstIII (α2,3-sialyltransferase)^11^. Both sialyltransferases belong to the, so far, monospecific CAZy (Carbohydrate-active enzymes database)^12–14^ glycosyltransferase (GT) family 42^15–17^. Although the presence of GT-42 and *N*-acetylneuraminate biosynthesis genes (*neu*ABC) is insufficient for expressing molecular mimics, all *C. jejuni* strains containing this set of genes^8,18–20^ (LOS locus classes A, B, C, M, and R) have been shown to synthesize ganglioside-like structures^3,7–9,21^. Therefore, the presence of GT-42 and *neu*ABC genes has been used as proxy for identifying *C. jejuni* strains capable of producing human glycan mimics^9,22^.

*Campylobacter coli* is the second most common cause of campylobacteriosis contributing, depending on the geographical region, to as many as 25% of all the infections^23^. Although *C. coli* has also been isolated from GBS patients^24,25^, its role in promoting this autoimmune disease remains controversial^26^. Additionally, despite the pervasive introgression with *C. jejuni*^27^, *C. coli* containing *C. jejuni-like* LOS classes linked to ganglioside mimicry have not been reported so far. Based on genomic data analysed hitherto, *C. coli* LOS locus appears to be marginally affected by horizontal gene transfer (HGT) or homologous recombination^28^.

Discovery of alternative orthologues of GT-42 encoding genes and associated LOS locus classes has been hindered by the very limited availability of genomic data. Consequently, it was only recently that *C. coli* LOS locus classes containing putative sialyltransferases, distantly related to those found in *C. jejuni*, were described^28,29^. The *C. coli* LOS locus class IX contains a GT-42 (*cstV*) and *neuABC* genes, LOS class II harbours an orphan GT-42 (*cstIV*), and LOS class III has a pseudogenized orphan GT-42^20,28,29^.

At present, the decreasing costs of next generation sequencing has driven a mass production of genomic sequences of several bacterial pathogens including *Campylobacter* spp. At the time of writing, the approximately 12,000 *C. jejuni* and 3,000 *C. coli* genome sequences found in public repositories offer unforeseeable opportunities. Thus, we took advantage of the large number of sequenced *Campylobacter spp*. strains to extract insights of significance for the debate of *C. coli* as an aetiological agent of GBS, aiming to comprehensively investigate presence, frequency, and distribution of the molecular machinery for the biosynthesis of sialylated LOS structures in *C. coli* population.

## MATERIAL AND METHODS

### Genome sequences mining, genes detection and allele calling

All whole genome raw sequence reads of entries deposited in the European Nucleotide Archive (ENA) as either *Campylobacter coli* or *Campylobacter jejuni* at the time of analysis (August 2017) were mapped against a set of reference genes (see below) for performing variant calling and inferring presence or absence using the ReMatCh framework v3.2 (<u>https://github.com/B-UMMI/ReMatCh</u>)^30^. Briefly, ReMatCh interacts with ENA for extracting and downloading all publicly available raw Illumina™ reads in *fastq* format for a given taxon. Then, it maps the reads onto the desired target loci using Bowtie2^31^, and performs variant calling with Samtools/Bcftools^32^ and ReMatCh Single Nucleotide Polymorphism call criteria. The minimum coverage depth to consider a position to be present in the alignment was fixed at 5 reads, and to perform allele calling the threshold was 10 reads. A locus was considered to be present if 1) at least 70% of the target reference gene sequence was successfully mapped and 2) if the consensus sequence was ≥ 80% identity at nucleotide level. When needed, the consensus sequence alignment was extracted using the script *combine_alignment_consensus.py* available in ReMatCh utilities.

### Identification and frequency of *C. coli* GT-42 homologues

To collect a set of *C. coli* reference genes homologous to *C. jejuni* GT-42 encoding genes, amino acid sequences of CstI (Uniprot Q9RGF1), CstII (Uniprot Q9F0M9) and CstIII (Uniprot Q7BP25) were used to search non-redundant (nr) NCBI protein sequences collection using blast+ V 2.7.1^33^ for best *C. coli* blastp hits (> 30% of amino acid identity; > 50% query coverage). Partial sequences were discarded and the remaining ones were used for an all-versus-all blastp analysis. Sequences were then categorized in separate groups having > 0.7 of Blastp Score Ratio (BSR)^34^. A Minimum Evolution phylogenetic tree based on the back-translated nucleotide sequence alignments (built with MUSCLE^35^ with default parameters) of all detected *C. coli* GT-42 proteins and *C. jejuni cstI, cstII*, and *cstIII* was inferred using MEGA7^36^. Finally, the detected *C. coli* GT-42 nucleotide sequences were used as reference for calling orthologues in all *C. coli* and *C. jejuni* strains using ReMatCh as described above.

### Identification of *Campylobacter coli* clades

To assign *C. coli* samples to one of the three previously described major phylogenetic clades^27,29^, population structure analysis and inferred phylogenetic relationships based on *atpA* gene^37^ were performed. The *atpA* sequence of *C. coli* strain RM2228 (KF855277) was used for allele calling in all *C. coli* strains using ReMatCh as described above. Based on the ReMatCh *atpA* consensus sequence alignment, samples were clustered using hierBAPS^38^ at first level and a Neighbor joining phylogenetic tree was inferred using MEGA7^36^. Representative strains from each *C. coli* clade^27,29^ were used as reference for classifying the clusters, and a set of *C. jejuni* strains were used as outgroup. The generated tree was visualized in iTOL^39^.

### Classification into LOS classes

To assign samples to one of the previously described *C. coli* LOS locus classes, nucleotide sequences of loci located between the “conserved putative two-domain glycosyltransferase” (orthologue 16 as described previously^20,28^) and the “LOS biosynthesis glycosyltransferase *waaV*” (orthologue 10 described previously^20,28^) from *C. coli* LOS locus classes I to XII^28^ were used for calling orthologues in all GT-42 positive *C. coli* using ReMatCh, as described above. Results were reported as percentage of genes present for a given LOS locus class.

### Pangenome analysis and gene flow investigation of LOS loci

For a set of *C. coli* strains of interest, raw sequencing data were retrieved from ENA with getSeqENA (<u>https://github.com/B-UMMI/getSeqENA</u>). Then, the paired-end raw reads were assembled using the INNUca pipeline (<u>https://github.com/INNUENDOCON/INNUca</u>), which consists of several modules and QA/QC steps. In brief, INNUca starts by calculating if the sample raw data fulfill the expected coverage (min 15x). After subjecting reads to quality analysis using FastQC (<u>http://www.bioinformatics.babraham.ac.uk/projects/fastqc/</u>), and cleaning with Trimmomatic^40^, INNUca proceeds to *de novo* draft genome assembly with SPAdes 3.11^41^ and checking assembly depth of coverage (min 30x). Finally, Pilon^42^ improves the draft genome by correcting bases, fixing misassembles, and filling gaps, prior species confirmation and MLST prediction with *mlst* software (<u>https://github.com/tseemann/mlst</u>).

Draft genomes passing INNUca QA/QC were annotated with Prokka^43^, and pangenome analysis was executed using Roary^44^ (default parameters). To annotate novel LOS locus classes, assemblies were manually inspected with Artemis^45^.

Horizontal Gene Transfer (HGT) among *C. coli* clades was inferred by mapping presence/absence of LOS associated group of orthologues into the *atpA* tree (see above). To infer possible gene transfer between *C. coli* and *C. jejuni*, representative sequences of LOS associated group of orthologues were blastn against nt NCBI database and HGT was detected if the best blast hit for *C. jejuni* was > 90% nucleotide identity over > 70% of the *C. coli* query length.

### Statistical analysis

Fisher's exact test was used to assess clade and GT-42 associations. P values of ≤0.05 were considered significant.

### Data availability

Data are available in Supplementary Information.

## RESULTS

### *C. coli* GT42 genes

Of the 45 *C. coli* GT42 protein sequences retrieved from NCBI nr database, six were partial sequences (i.e. incomplete coding sequences). Thus, they were excluded from further analysis (Supplementary Table S1). Based on BSR, the remaining 39 sequences clustered into 7 different groups (Supplementary Table S2 and S3), with average BSR values ranging from 0.80 to 0.98 (Table 1). Group 1, 2, and 4 contain proteins showing the highest similarity to CstI, CstIII, and CstII, respectively, while the other groups show limited homology to *C. jejuni* GT-42 enzymes (Table 1). Group 5 comprises orthologues to the previously described CstV in LOS class IX of *C. coli* 76339^29^, while Group 7 includes CstIV, the GT-42 within *C. coli* LOS locus class II ^20,28^. Group 6 contains a novel group of orthologous proteins (named herein CstVI) showing high similarity to the pseudogenizised GT-42 described as part of LOS locus class III^20,28^. Similarly, Group 3 includes a single novel protein sequence named herein CstVII.

**Table 1.** Average Blastp Score Ratio (BSR) of the *C. coli* GT-42 homologs

| Groups | Gene<sup>ref</sup> | BSR | CstI (Q9RGF1) |  | CstII (Q9F0M9) |  | CstIII (Q7BP25) |  |
| --- | --- | --- | --- | --- | --- | --- | --- | --- |
|  |  |  | Cov (%) | Id (%) | Cov (%) | Id (%) | Cov (%) | Id (%) |
| 1 | *cstI*<sup>21</sup> | 0.87 | 93 | 70 | 89 | 51 | 83 | 50 |
| 2 | *cstIII*<sup>21</sup> | 0.80 | 56 | 54 | 98 | 51 | 100 | 88 |
| 3<sup>a</sup> | *cstVII*<sup>c</sup> | 1 | 61 | 51 | 90 | 52 | 86 | 53 |
| 4 | *cstII*<sup>21</sup> | 0.87 | 60 | 52 | 99 | 89 | 95 | 52 |
| 5 | *cstV*<sup>29</sup> | 0.98 | -<sup>b</sup> | - | 98 | 48 | 94 | 43 |
| 6 | *cstVI*<sup>c</sup> | 0.95 | - | - | 97 | 37 | 96 | 35 |
| 7 | *cstIV*<sup>29</sup> | 0.86 | - | - | 97 | 40 | 94 | 37 |

<sup>a</sup>singleton
<sup>b</sup>no significant hits
<sup>c</sup>gene name proposed in this study

Furthermore, evolutionary analysis revealed that the 7 BSR groups form monophyletic clades and are divided into two clusters (Fig. 1). Cluster A is comprised of *cstI, cstII, cstIII*, and *cstVII*, while cluster B includes *cstIV, cstV* and *cstVI*.

**Figure 1.**
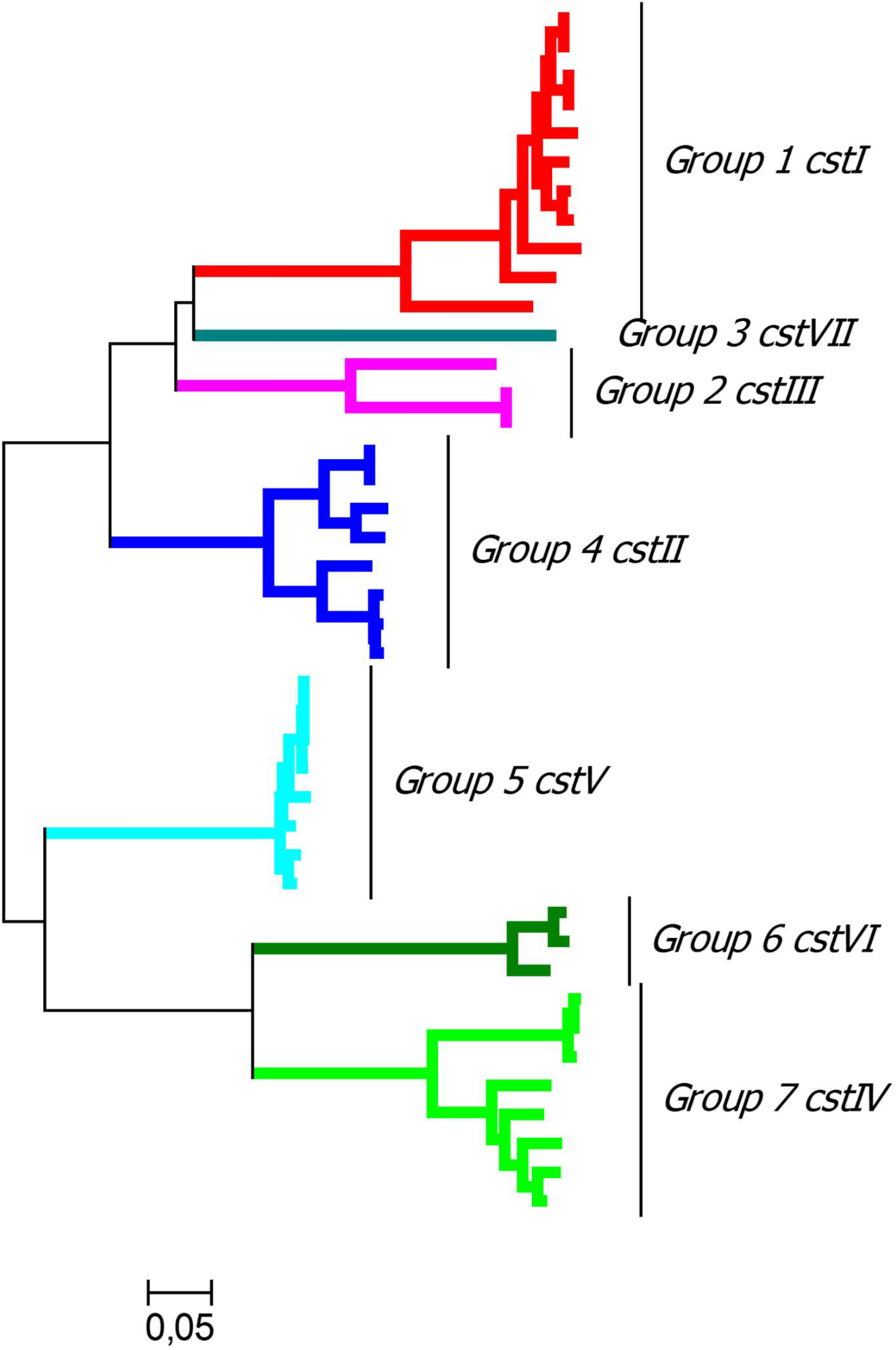
Evolutionary analyses of *C. coli* GT-42. Evolutionary analysis of 45 *C. coli* GT-42 sequences and three *C. jejuni* sequences (*cstI, cstII* and *cstIII*) was conducted in MEGA7 and the evolutionary history was inferred using the Minimum Evolution method calculating the distance using Maximum Composite Likelihood. The tree is drawn to scale, with branch lengths in the same units (number of base substitutions per site) as those of the evolutionary distances used to infer the phylogenetic tree.

### Prevalence of GT-42 encoding genes in *C. coli* population

Raw reads from 2,582 genomes submitted as *C. coli* were retrieved from ENA and classified into one of the three major *C. coli* phylogenetic clades based on *atpA* phylogeny and hierBAPS clustering (Supplementary Fig. S1). A total of 29 genomes were excluded from further analyses, as *atpA* phylogenetic analysis confirmed them to be *C. jejuni*. Altogether, 2,432 (95%) genomes belonging to Clade 1, 40 (1.6%) to Clade 2 and 81 (3.2%) to Clade 3 were mapped against all the sequences classified into the 7 *C. coli* GT-42 groups. A total of 818 (32%) *C. coli* genomes were positive for at least one GT-42 encoding gene (Table 2; Supplementary Table S4). GT-42 genes were found in approximately one third of *C. coli* Clade 1 (774/2,432; 31.8%). Furthermore, GT-42 genes were underrepresented in *C. coli* Clade 2 (2/40; 5%; P < 0.0001), while overrepresented in Clade 3 (42/81; 52%; P < 0.001). Overall, cluster B GT-42 genes (*cstIV, cstV* and *cstVI*) were the most abundant GT-42 detected in the *C. coli* population, accounting for 84.2% of the alleles. Conversely, cluster A GT-42 genes (*cstI, cstII, cstIII* and *cstVII*) only represented 15.8% of the alleles (Table 2). The most abundant *C. coli* GT-42 was *cstVI*, whereas *cstIII* was the rarest. *C. coli* Clade 1 strains were overrepresented in *cstVII* and *cstVI*, and underrepresented in *cstV* (P < 0.01). Conversely, Clade 3 strains were underrepresented in *cstVII* and *cstVI*, and overrepresented in *cstV* (P < 0.01).

**Table 2.** Distribution of GT-42 genes among *C. coli* clades.

| *C. coli* | GT-42 Cluster A |  |  |  | GT-42 Cluster B |  |  | Total |
| --- | --- | --- | --- | --- | --- | --- | --- | --- |
|  | *cstI*<br>(G1) | *cstII*<br>(G4) | *cstIII*<br>(G2) | *cstVII*<br>(G3) | *cstIV*<br>(G7) | *cstV*<br>(G5) | *cstVI*<br>(G6) |  |
| Clade 1 | 2 | 22 | 2 | 91 | 267 | 0 | 414 | 798 (94%) |
| Clade 2 | 0 | 0 | 2 | 0 | 0 | 0 | 0 | 2 (0.2%) |
| Clade 3 | 10 | 5 | 0 | 0 | 14 | 15 | 5 | 49 (5.8%) |
| Total | 12<br>(1.41%) | 27<br>(3.19%) | 4<br>(0.47%) | 91<br>(10.73%) | 281<br>(33.10%) | 15<br>(1.77%) | 419<br>(49.35%) | 849<br>(100%) |

### *C. coli* LOS classes contain GT-42 encoding genes

To predict the LOS locus composition of GT-42 positive strains, genomes were mapped against all genes from all known LOS locus classes^28^. Results are available in Supplementary Table S4. The presence of GT-42 gene alleles from cluster B was strongly concordant with predicted LOS locus classes. For *cstVI* positive strains, 99% were predicted to have a LOS locus class III-like. Similarly, 93% of *cstV* positive *C. coli* possessed a LOS locus class IX-like, and 68% of *cstIV* positive strains harboured a LOS locus class II-like. Contrastingly, genomes exclusively positive for cluster A GT-42 genes had no significant match to any of the previously defined LOS locus classes (Supplementary Table S4).

To determine the exact genetic composition and synteny of the LOS loci, 261 GT-42 positive genomes were assembled and manually inspected. The data set included all Clade 2 and 3 strains and a selection of Clade 1 strains comprising all *cstI, cstII, cstIII*, and *cstVII* positive strains, and a subset of randomly selected *cstIV* and *cstVI* positive strains (Supplementary Table S4). Annotation of the identified LOS locus classes is available in Supplementary Table S5. Apart from *cstI* and *cstVII*, all GT-42 genes were found within the LOS locus. Among assembled genomes, 61.3% (160) were found to contain a LOS-associated GT-42 gene. Besides the three previously described LOS locus classes containing GT-42 genes (i.e. classes II, III, and IX), 23 novel classes were identified (Fig. 2). LOS class III was the most abundant accounting for 72 isolates, followed by II (39), XXIII (9), XXV (8), XXVI (5), XXX (4), XVI (2), XXVIII (2), and XXXIII (2). The rest of the classes (17) were represented by a single strain. A strong association between LOS locus composition, *C. coli* Clade, and GT-42 gene alleles, was observed. In general, *C. coli* Clade 1 exhibited lower LOS locus diversity compared to the other clades. In Clade 1, genomes positive for *cstIV* and *cstVI* (88.9% of the total) possess LOS locus classes II and III, respectivelly, with 99% nt sequence identity. In all cases *cstVI* was present as a pseudogene. Contrastingly, Clade 3 *C. coli* evince a larger genetic variability in LOS locus classes containing *cstIV* (8 classes), *cstV* (3), or *cstVI* (2). Interestingly, no pseudogenes were found.

Albeit the rarity of LOS associated cluster A GT-42 genes (*cstII* and *cstIII*) in *C. coli* population (1.2%), several distinct LOS locus classes were identified (Fig. 2a). Out of the ten LOS classes containing *cstII* (Fig. 2a), only XXIII was detected in multiple Clade 1 (7) and Clade 3 (3) strains. Meanwhile, *cstIII* was located in two different LOS locus classes in *C. coli* Clade 2 strains.

**Figure 2.**
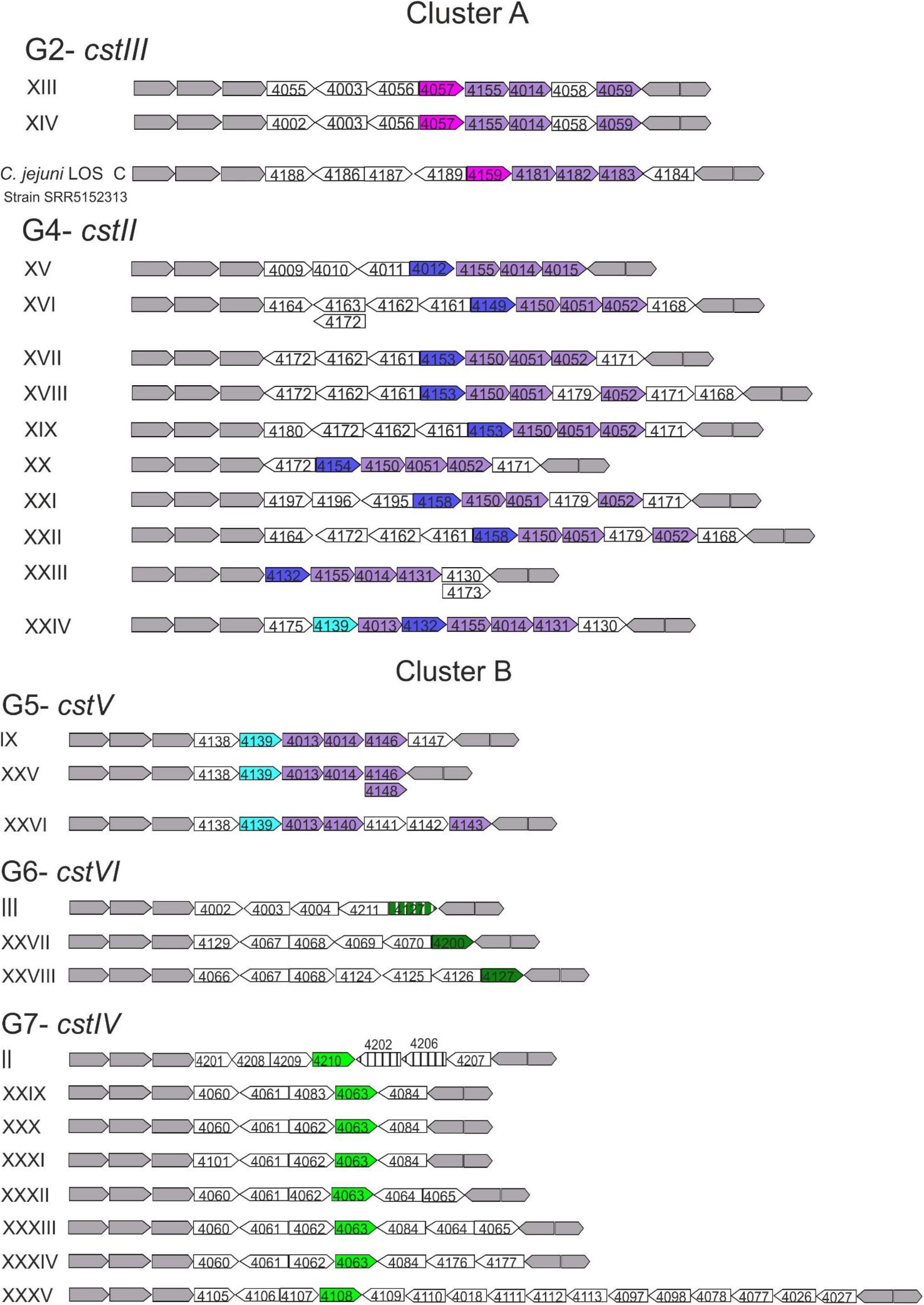
*C. coli* LOS classes containing GT-42 genes. Arrows represent open reading frames. White arrows: genes putatively unrelated to biosynthesis and transfer of Neu5Ac. Grey arrows: conserved genes. Purple arrows: sialic acid biosynthesis genes always present *neuB, neuC*, and *neuA* order. Pink arrows: GT-42 orthologues from group 2. Dark blue arrows: GT-42 orthologues from group 4. Light blue arrows: GT-42 orthologues from group 5. Dark green arrows: GT-42 orthologues from group 6. Light green arrows: GT-42 orthologues from group 7. Striped genes are fragmented. Representation of LOS class II was adapted to reflect origin from LOS class XXXIV. Gene size is not drawn to scale.

All LOS locus classes containing *cstII, cstIII*, or *cstV* were positive for *neuABC* genes. Contrastingly, only 6.44% and 4.51% of *cstIV* and *cstVI* positive strains, respectively, contained *neuABC* genes which were invariably located outside the LOS locus and frequently in association with *cstI* or *cstVII*.

### Gene flow and evolution of GT-42 containing LOS locus classes in *C. coli*

Based on orthologue group delineation by Roary (>95% amino acid identity), strains belonging to different *C. coli* clades were shown to share LOS-associated orthologues (Fig. 3). Hence, suggesting gene flow of LOS genes across *C. coli* clades. Interestingly, most of the share orthologues between clades encode proteins putatively involved in sugar biosynthesis or sugar modification (Table 3). Insights into the evolution of *C. coli* GT-42 containing LOS classes were gain by comparing Clade 1 with Clade 3 LOS classes. Reciprocal blastn analysis between LOS locus classes II (Clade 1) and class XXXIV (Clade 3) showed ~88% nucleotide identity over ~99% of length. Likewise, the terminal part of LOS class III showed high similarity (>90% nucleotide identity) to LOS classes XXVII and XXVIII. Notably, in both Clade 1 LOS locus classes gene pseudogenization was observed: the phosphoethanolamine transferase genes (*eptC*) in class II, and *cstVI* in class III. Thus, LOS locus classes II and III plausibly originated from Clade 3 LOS classes and underwent a diversification process (including pseudogenization) and clonally expanded as a consequence of adaptation to the agricultural niche.

**Figure 3.**
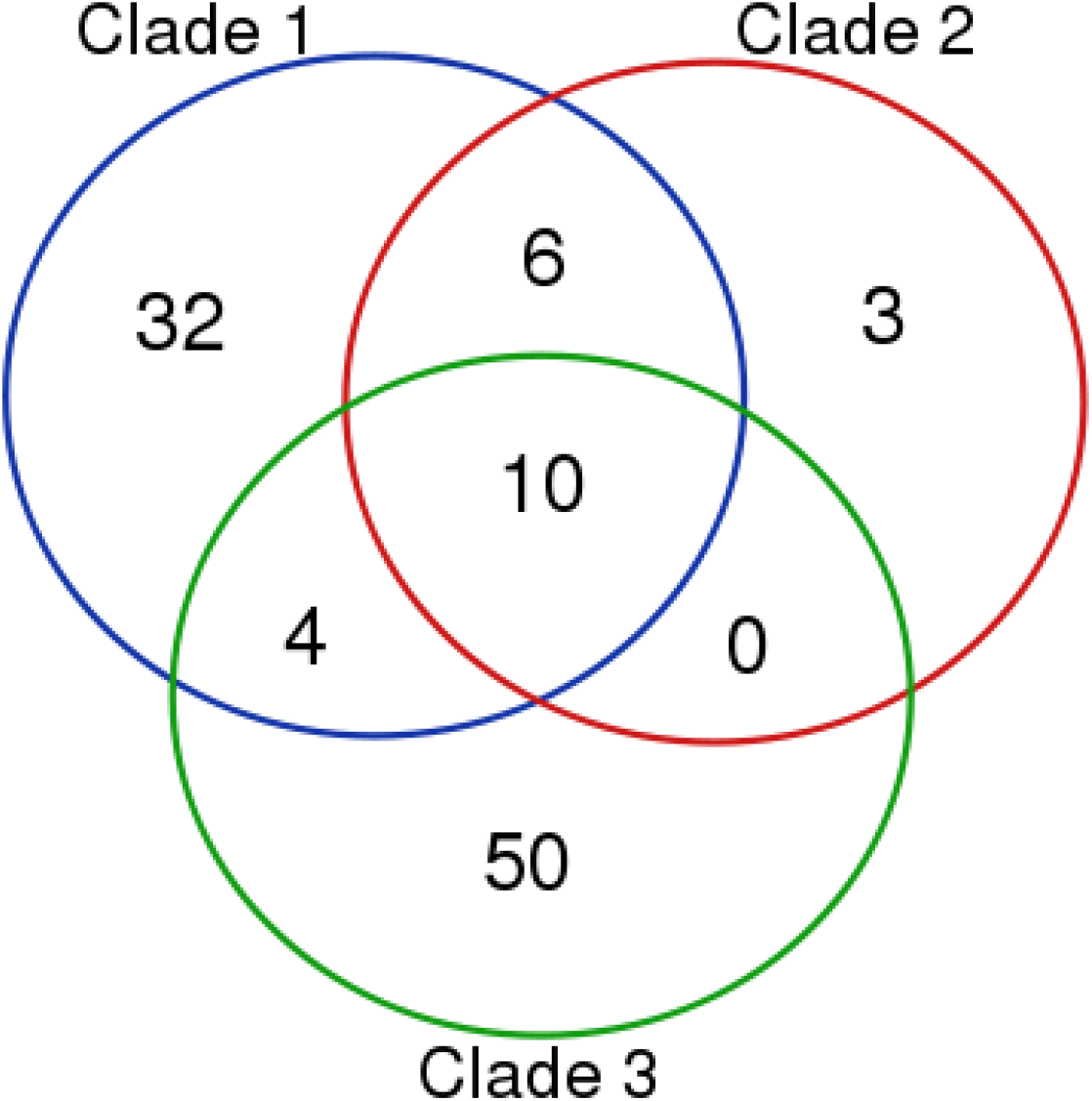
Venn diagram illustrating the number of orthologues shared between *C. coli* major clades. 10 orthologue were found in all three *C. coli* clades.

**Table 3.** Group of orthologues shared among *C. coli* clades

| Roary Orthologue | Prokka annotation | 1 | 2 | 3 |
| --- | --- | --- | --- | --- |
| 4026 | dTDP-glucose 4,6-dehydratase | + | + | + |
| 4027 | Glucose-1-phosphate thymidyltransferase | + | + | + |
| 4078 | TDP-4-oxo-6-deoxy-alpha-D-glucose-3,4-oxoisomerase | + | + | + |
| 4014 | UDP-N-acetylglucosamine 2-epimerase | + | + | + |
| **4155** | N,N'-diacetyllegionaminic acid synthase | + | + | + |
| **4077** | Polysialic acid O-acetyltransferase | + | + | + |
| **4042** | UDP-glucose 6-dehydrogenase | + | + | + |
| **4041** | UDP-glucose 4-epimerase | + | + | + |
| **4076** | UDP-glucose 4-epimerase | + | + | + |
| **4075** | UDP-galactopyranose mutase | + | + | + |
| **4003** | General stress protein A | + | + | - |
| **4002** | GalNAc-alpha-(1->4)-GalNAc-alpha-(1->3)-diNAcBac-PP-undecaprenol alpha-1,4-N-acetyl-D-galactosaminyltransferase | + | + | - |
| **4004** | putative glycosyltransferase EpsJ | + | + | - |
| **4051** | putative glycosyltransferase EpsJ | + | + | - |
| **4059** | N-acylneuraminate cytidyltransferase | + | + | - |
| **4052** | N-acylneuraminate cytidyltransferase | + | + | - |
| **4127** | hypothetical protein | + | - | + |
| **4126** | hypothetical protein | + | - | + |
| **4132** | hypothetical protein | + | - | + |
| **4157** | hypothetical protein | + | - | + |

### Prevalence of GT-42 genes in *C. jejuni*

Prevalence of GT-42 homologues in *C. jejuni* was investigated by mapping 12,391 genome sequences deposited as *C. jejuni* against the 7 *C. coli* GT-42 groups. A total of 61.15% of the putative *C. jejuni* genomes were positive for at least one gene. Unsurprisingly, *cstII* and *cstIII* were the most abundant representing 95.75% of the GT-42 sequences detected. The remaining gene groups were either present in a minority of the tested genomes (*cstIV*, 101; *cstVI* 211; *cstVII*, 2) or non-detected (*cstV*). Genomes positive to GT-42 sequences other than *cstII* and *cstIII* were assembled for species verification and to manually inspect the LOS locus gene composition. Only 52 (16.6%) genomes were confirmed as *C. jejuni* by INNUca (Supplementary Table S6), 40 of which (77%) were positive for *cstIV*, 10 to *cstVI* (19.2%), and 2 to *cstVII* (3.8%). Similarly to *C. coli, C. jejuni cstVII* was located outside the LOS locus and downstream from *neuABC* genes.

### Introgression between *C. jejuni* and *C. coli* affect GT-42 containing LOS classes

High similarity between *C. coli* and *C. jejuni* cstII-associated LOS locus was observed (i.e. >80% gene lengths and >95% of nucleotide identity), implying recent gene flow between the two species. In fact, *C. coli* LOS classes XVI, XVII, XVIII, XIX, XX, XXI, and XXII are a mosaic of *C. jejuni* LOS classes A, B, S/F, and I/D. *C. coli* LOS class XXIV is further evidence of admixture between the two species, as it includes the *C. coli* specific *cstV* and *neuB* orthologues, as well as the *C. jejuni cstII* and *neuB* copies (Supplementary Table S4, Fig. 2). Finally, cross-species mobilization of an entire LOS locus classes was also encountered. *C. coli* strain SRR5152313 carries *C. jejuni* 11168 LOS class C, and 35 out of 40 *cstIV* positive *C. jejuni* strains, 37.1% of which from MLST sequence type 459, harbour a *C. coli* LOS locus class II.

## DISCUSSION

The small number of GBS associated *C. coli* isolates and the supposedly absence of molecular machinery for ganglioside mimicry are the main reasons, so far, supporting the idea of no link between *C. coli* and GBS. In 1994 von Wulffen and colleagues reported the first *C. coli* isolated from a GBS patient in a comparative seroreactivity study^46^. The *C. coli* strain in question exhibited a Lior type 11 phenotype, which had also been found in GBS-associated *C. jejuni* strains. Thus, in the following years *C. coli* was considered as a plausible GBS causing organism^46^. However, after recognition of *C. jejuni* expressing ganglioside-like LOS as the infectious agent triggering GBS, testing for cross-reactivity with anti-ganglioside autoantibodies became critical in understanding GBS aetiology. So far, insufficient evidence supporting a causal relationship between *C. coli* and GBS has been found since studied GBS-associated *C. coli* strains have been unreactive to monoclonal anti-ganglioside antibodies^24,25^. Furthermore, although the GBS-associated *C. coli* strain 664H2004 has been shown to carry a *cstII* orthologue and di-sialylated LOS, no further evidence suggesting expression of ganglioside mimics was attained, as authors failed to genetically and structurally characterize *C. coli* 664H2004 LOS^26^.

In the present study, 16 *C. coli* LOS locus classes (Fig. 2) were shown to contain the essential molecular machinery to potentially express sialylated LOS (i.e. a *cst* homologue and *neuABC*). While genotype is generally insufficient to predict LOS structure^3,8,19^, considerable evidence supporting the expression of ganglioside-like LOS in *C. coli* was found. In contrast to previous reports^27,28^, *C. coli* LOS locus may be substantially affected by introgression with *C. jejuni*. Herein, 10 *C. coli* LOS locus classes containing a *cstII* were demonstrated to be mosaics of *C. jejuni* LOS classes. *C. jejuni* strains carrying *cstII* containing LOS classes have hitherto rarely being found to express non-ganglioside sialylated LOS^3,8,9,18,21^. Furthermore, extreme introgression resulted in acquisition of the entire *C. jejuni* LOS class C in *C. coli* SRR5152313 (100% homology). Consequently, this strain, isolated from turkey in US in 2016, could potentially trigger GBS as most likely expresses a GM1a- or GM2-like LOS ^9^.

However, it is to be noted that strains carrying *C. jejuni*-like LOS locus are a minority in the *C. coli* population (approximately 1% of sequenced strains). Most of the *C. coli* possessing GT-42 genes (i.e. *cstIV* and *cstVI*) carry LOS classes lacking *neuABC* genes (approximately 27% of the sequenced strains). Furthermore, genome-wise analysis failed to identify genes potentially linked to the synthesis of CstIV and CstVI sugar donors. Even though functional studies are needed to clarify the activity of CstIV and CstVI, it seems plausible to believe that these elements are not involved in LOS ganglioside mimicry based on the results presented here and the absence of Neu5Ac in the LOS of *cstIV* positive strains^28^. Thus, the infrequency of the genetic structures related to ganglioside mimicry in the population might be the reason behind *C. coli* little contribution to GBS incidence^26^.

Beside ganglioside mimicry and the pathogenesis of GBS, expression of sialylated structures has a strong impact on host-bacteria interaction^10^. In our broad-gauge screening, we have shown that a considerable proportion of *C. coli* strains carry GT-42 genes within the LOS locus (29% of *C. coli* deposited in ENA at the time of writing). Overall, 23 new GT-42 associated LOS classes were described, 15 of which were present exclusively in the non-agriculture *C. coli* belonging to Clade 3.

Thus, underrepresentation of non-agricultural *C. coli* strains^27^ in studies characterizing the LOS loci of extensive strain collections^20,28,29^ probably hampered earlier identification of a wider diversity of LOS classes with GT-42 genes.

We also discovered that LOS locus classes II and III^28^, the most predominant among agriculture-adapted Clade 1 *C. coli*, most likely originated from non-agriculture Clade 3 LOS classes. Moreover, few genes in both classes, including the GT-42 *cstVI*, lost their function in Clade 1. Cell surface structural changes as result of natural selection is a dominant phenomenon in microbial evolution. In pneumococcus, for example, natural selection as a consequence of vaccination programs targeting polysaccharide structures has resulted in shifts in the population of nonvaccine-type strains^47^. Outer membrane or wall-associated structures in bacteria (i.e. oligo and polysaccharides and proteins) play also a fundamental role in host interaction. Thus, they are subjected to diversifying selective pressure to conform to distinct receptors in different host species^48^. Moreover, reductive evolution leading to functional loss of several genes through e.g. pseudogenization is a common feature of bacterial undergoing niche adaptation^48^. For example, a single naturally occurring nucleotide mutation responsible for the inactivation of a gene essential for D-alanylation of teichoic acids, has been shown to be sufficient to convert a human-specific *Staphylococcus aureus* strain into one that could infect rabbits^49^. Introduction of the agricultural niche was key in the evolution of *C. coli* clades^50^: clade 1 expanded within this niche and underwent an extensive genome introgression with *C. jejuni^27^*.

Therefore, it is tempting to speculate that gene loss within imported LOS classes II and III, may have played a significant role in the expansion of *C. coli* in the agricultural niche by shaping the outer membrane composition. This hypothesis is supported by two pieces of evidence: (i) the predominance of LOS classes II and III in *C. coli* Clade 1 generalist (i.e. multihost) strains^28^ and (ii) the strong purifying selection resulting in limited nucleotide variability in these LOS locus classes (>99% identity. The importance of LOS locus classes II and III in adaptation to the agricultural niche is further evidenced by the flow of these genetic elements between *C. coli* and agricultural *C. jejuni*. Although introgression between *C. jejuni* and *C. coli* has been considered to be unilateral until now^27^, we identified several *C. jejuni* strains carrying LOS classes typically detected in *C. coli* Clade 1. Most of the *C. jejuni* strains carried a LOS class II with 99% identity, while some other presented a mosaic of Clade 3 LOS classes containing *cstIV*. As described previously^22^, a strong association between MLST type and LOS class was observed, being the bovine associated ST-459^51^ the most prevalent among the *C. jejuni* carrying *C. coli* LOS class II.

## CONCLUSION

Although at extremely low frequencies, bacterial factors implicated in GBS aetiology can cross clade and species barriers. Furthermore, spreading of these factors in the population could potentially result in *C. coli* playing a more prominent role in GBS. *C. coli* also presents a larger GT-42 enzyme repertoire than *C. jejuni*. Nevertheless, the activity of these enzymes and their role shaping *C. coli* population is yet to be explored. Overall, *C. coli* glycobiology is largely unknown in spite of being a major foodborne pathogen.

## ACKNOWLEDGMENTS

This study was funded by the following grants; University of Helsinki three years research grant 313/51/2013, ONEIDA project (LISBOA-01-0145-FEDER-016417) co-funded by FEEI - “Fundos Europeus Estruturais e de Investimento” from “Programa Operacional Regional Lisboa 2020” and by national funds from FCT - “Fundação para a Ciência e a Tecnologia” and BacGenTrack (TUBITAK/0004/2014) [FCT/ Scientific and Technological Research Council of Turkey (Türkiye Bilimsel ve Teknolojik Araşrrma Kurumu, TÜBITAK)]”. A. C was supported by the Microbiology and Biotechnology graduate program from the University of Helsinki. The authors wish to thank CSC- Tieteen tietotekniikan keskus Oy for providing access to cloud computing resources.

## CONTRIBUTIONS

A.C designed and coordinated the study. A.C and M.R performed data analysis, prepared figures, and wrote the manuscript. J.A.C. and M.P.M design and developed INNUca and ReMatCh. All authors have contributed to data interpretation, have critically reviewed the manuscript, and approved the final version as submitted.

